# Lytic transglycosylases RlpA and MltC assist in *Vibrio cholerae* daughter cell separation

**DOI:** 10.1101/608497

**Authors:** Anna I. Weaver, Valeria Jiménez-Ruiz, Srikar R. Tallavajhalla, Brett P. Ransegnola, Kimberly Q. Wong, Tobias Dörr

## Abstract

The cell wall is a crucial structural feature in the vast majority of bacteria and comprises a rigid, covalently closed, mesh-like network of peptidoglycan (PG) strands. While PG synthesis is important for bacterial survival under many conditions, the cell wall is also a dynamic structure, undergoing degradation and remodeling by so-called “autolysins”, enzymes that break bonds in the PG network. Cell division, for example, requires extensive PG remodeling and separation of daughter cells, which depends primarily upon the activity of amidases. However, in *V. cholerae*, we have found that amidase activity alone is insufficient for daughter cell separation and that the lytic transglycosylases RlpA and MltC both contribute to this process. MltC and RlpA both localize to the septum and are functionally redundant under normal laboratory conditions; however, only RlpA can support normal cell separation in low salt media. The division-specific activity of lytic transglycosylases has implications for the local structure of septal PG, suggesting that there may be glycan bridges between daughter cells that cannot be resolved by amidases. We propose that lytic transglycosylases at the septum serve as a back-up mechanism to cleave rare, stochastically produced PG strands that are crosslinked beyond the reach of the highly spatio-temporally limited activity of the amidase and to clear PG debris that may block the completion of outer-membrane invagination.

## INTRODUCTION

The cell wall is a crucial structural feature for the vast majority of bacteria and is mainly composed of a rigid, yet elastic, covalently-bound network of peptidoglycan (PG) strands. PG has an oligomeric glycan backbone that is assembled by glycosyltransferases (GTs, RodA/FtsW and class A Penicillin Binding Proteins [aPBPs]) (Cho et al., 2016; Leclercq et al., 2017; Zhao et al., 2017; Taguchi et al., 2019) through the polymerization of *N*-acetylglucosamine (NAG) and *N*-acetylmuramic acid (NAM) heterodimers. These PG strands are crosslinked to adjacent strands primarily by the transpeptidase domains of aPBPs and bPBPs via short peptides attached to the NAM residues, resulting in the strong, mesh-like sacculus. While the rigidity functions to resist bacterial cells’ high internal pressure (Osawa & Erickson, 2018), it restricts processes such as cell growth, division, and insertion of multiprotein trans-envelope complexes such as the flagellum and secretion systems (Nambu et al., 1999; Santin & Cascales, 2017). The cell wall must therefore be a dynamic structure, and indeed undergoes constant remodeling and recycling by PG degradation enzymes collectively known as “autolysins” (T. K. Lee & Huang, 2013).

Autolysins are numerous and diverse, in part owing to the complexity of the substrate on which they act. Almost all different types of covalent bonds that are found within PG can be cleaved by autolysins, and many of these enzymes are functionally redundant under standard laboratory growth conditions. Functional redundancy has stymied the elucidation of the physiological role of many autolysins, as it makes them inaccessible to many traditional means of assessing gene-phenotype associations, such as analysis of single gene knockouts. The lytic transglycosylases (LTGs), for example, have been exceptionally well-characterized biochemically (Dik et al., 2017), but still relatively little is known about their individual physiological functions. LTGs target PG at the glycosidic bond between NAG and NAM residues and the primary mechanism for this cleavage is a non-hydrolytic, intramolecular cyclization of NAM to form 1,6-anhydroMurNac (^anh^NAM) (Holtje et al., 1975; Dik et al., 2017; Williams et al., 2018). At least in well-understood model organisms, this signature anhydro “cap” is assumed to be at the end of almost all peptidoglycan strands *in vivo* (Kraft et al., 1998; Heidrich et al., 2002a). Members of the LTG class have been implicated in many cellular processes, including the termination of GT-mediated PG polymerization (Tsui et al., 2016; Yunck et al., 2016), insertion of secretory apparati and flagella (Herlihey & Clarke, 2017; Santin & Cascales, 2017), pathogenesis (Chan et al., 2012), and PG recycling (Cloud & Dillard, 2002).

One process where PG remodeling is particularly important is cell division. Our current understanding of bacterial cell division includes a step in which lateral PG must be remodeled to allow for insertion of a septal wall between daughter cells, followed by cleavage of that septal wall to facilitate daughter cell separation (Potluri et al., 2012; Egan & Vollmer, 2013). Septal PG cleavage by amidases, which cleave off the dipeptide side stem from the NAM residue, is tightly controlled spatiotemporally to ensure that PG degradation is exclusively localized to where it is needed (Heidrich et al., 2002a; Uehara et al., 2010; D. C. Yang et al., 2011; Peters et al., 2011; Möll et al., 2014; Tsang et al., 2017). The amidases are generally assumed to be the main enzymes mediating daughter cell separation, though there is evidence that other autolysins, including LTGs, are pleiotropically involved (Heidrich et al., 2001, 2002a; Priyadarshini et al., 2006). *E. coli* strains lacking LTGs MltABCDE and Slt70, for example, have mild cell separation defects (Heidrich et al., 2002a). In addition, Jorgenson, et. al. identified a highly conserved LTG, RlpA, which exhibits septum-specific cleavage activity in *Pseudomonas aeruginosa.* RlpA is required under low salt conditions, but not during growth in standard laboratory media, suggesting *P. aeruginosa* encodes at least one redundant septal LTG (Jorgenson et al., 2014). *Salmonella enterica* similarly appears to require LTGs MltC and MltE for proper cell daughter cell separation in low salt conditions (Wall et al., 2011) and in *Neisseria gonorrhea*, mutating the LTG LtgC results in daughter cell separation defects (Cloud & Dillard, 2004). Thus, LTGs appear to play important and often redundant, but poorly-understood roles in septal cleavage in diverse bacteria. In particular, it is unclear whether the separation defects observed in LTG mutants are due to the lack of a septum-specific function of these autolysins, or a general consequence of pleiotropic cell wall damage.

Here we show that in the cholera pathogen, *Vibrio cholerae*, the two LTGs MltC and RlpA are collectively required for daughter cell separation. Their inactivation results in the generation of cell chains reminiscent of amidase mutants and additional deletion of *V. cholerae*’s sole amidase exacerbates this chaining defect. Our data suggest that these LTGs fulfill specialized roles in daughter cell separation and have important implications for septal PG architecture in the cholera pathogen.

## RESULTS

### Simultaneous inactivation of seven LTGs induces a lethal cell separation defect

Autolysins are often functionally redundant. *V. cholerae*, for example, can tolerate the inactivation of its sole amidase or simultaneous deletion of 5 out of its 6 M23 family endopeptidases (Dörr et al., 2013; Dörr, Davis, et al., 2015), suggesting that redundant autolysins can substitute for each other to sustain at least basic growth processes. In addition to the amidase and the endopeptidases, *V. cholerae*’s genome encodes 8 predicted LTGs that were identified based on their homology to *E. coli* LTGs. Whether these enzymes fulfill unique or redundant roles within *V. cholerae’*s life cycle is unknown. To find new phenotypes associated with LTG deficiency, we endeavored to make sequential deletions in all eight genes. Interestingly, we were able to inactivate six LTGs (*mltA, mltB, mltC, mltD, mltF* and *slt70*, leaving only *rlpA* and *mltG* intact) with the resulting strain exhibiting only slight morphological aberrations, such as a mild division defect with a corresponding increase in cell length (**Fig. 1A**, **Fig. S1A**). Thus, in *V. cholerae*, RlpA and MltG are cumulatively capable of performing all potentially essential functions of LTGs, at least under laboratory conditions.

**Fig 1.**
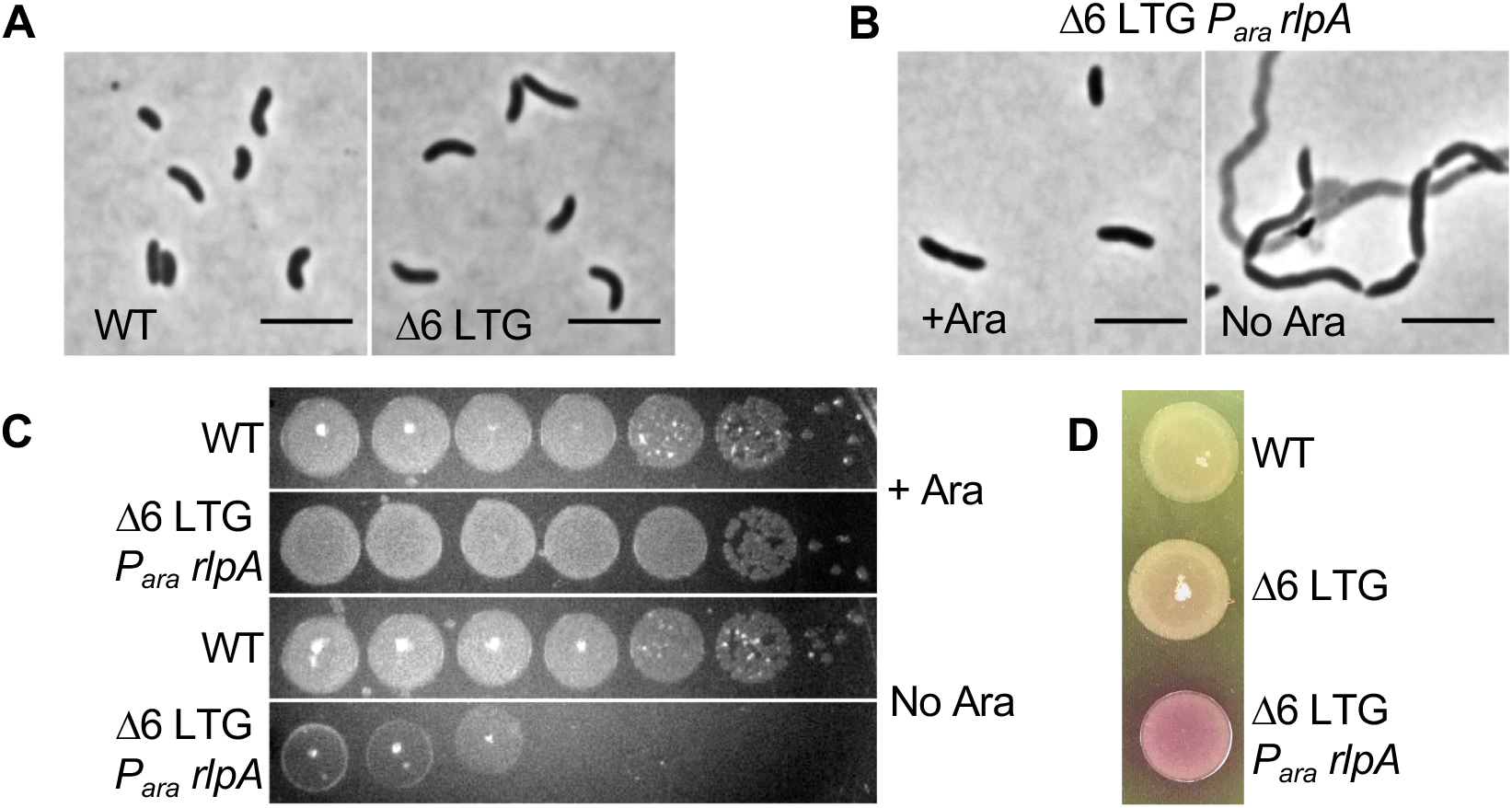
Lytic transglycosylases are collectively required for survival of *V. cholerae*. **A)** WT and Δ6 LTG cultures grown LB at 37°C and imaged on an agarose pad. **B)** RlpA was depleted in the Δ6 LTG background by placing its native promoter under control of arabinose induction and growing in the absence or presence of arabinose (ara). Cells were imaged on an agarose pad and **C)** spot-plated onto LB+/-0.2% ara, followed by incubation at 30°C for 24hrs. Grid lines = 1cm. **D)** Lysis was visualized by culturing strains overnight in LB + ara at 37°C, spotting 10µL directly onto an LB+20 µg mL^-1^ CPRG plate, and incubating 18hrs at 30°C. Scale bars = 5µm. All experiments are representatives of at least two biological replicates.

Though single knockouts of *mltG* and *rlpA* could be readily obtained in a wild-type background, (**Fig. S1B**) we were unable to further delete *mltG* or *rlpA* from the Δ6 LTG strain, suggesting that these two represent a minimal set of LTG functions required for viability. To test the phenotypic consequences of loss of RlpA in the Δ6 LTG background, we constructed a strain that expressed its sole copy of *rlpA* under control of an arabinose-inducible promoter. Growing this strain in the absence of arabinose resulted in the formation of long chains of unseparated cells, many of which had lysed (**Fig. 1B**). This lysis was also evident in the ∼10^6^-fold reduction in plating efficiency after RlpA depletion and dramatic color change when plated on the cell-impermeable β-galactosidase substrate, chlorophenol red-β-D-galactopyranoside (CPRG) (**Fig. 1CD**), an established readout of cell lysis (Paradis-bleau et al., 2014). The lethal chaining defect seen in the Δ6 LTG strain depleted of RlpA suggests that at least one of the roles of LTGs is an essential function related to septal PG remodeling and/or daughter cell separation that cannot be adequately fulfilled by other autolysins under native conditions. Since a similar depletion strain for MltG did not exhibit a phenotype, we here focused on RlpA for further study; the reason for MltG’s essentiality in the Δ6 background is subject of future work.

### Simultaneous inactivation of RlpA and MltC results in a cell separation defect

Since cell separation defects are not lethal in other *V. cholerae* autolysin mutants (*e.g.*, amidase mutants [Möll et al., 2014]), we hypothesized that the lethal phenotype of the Δ6 LTG/*rlpA* depletion strain was in principle separable from the chaining defect. To dissect this further, we assayed different combinations of LTG mutants for cell morphology defects **(Fig. S2A).** LTGs were generally inactivated by replacing their open reading frame with a scar sequence. However, the gene for RlpA is located within a genomic region containing other important cell wall factors, including *rodA* and *dacA*, and so we instead inserted a premature stop codon in position 133 to reduce potential polar effects. Visual inspection of different mutant combinations revealed that a strain deleted in both *mltC* and *rlpA* (Δ2 LTG) exhibited a pronounced cell separation defect, manifest as long chains in a culture grown to ∼OD_600_ 0.8 (**Fig 2A**). Imaging these chains expressing cytoplasmic GFP from a constitutive promoter revealed clearly separated cytoplasmic spaces, indicating that cytokinesis was complete in cell chains. In contrast to the Δ2 LTG chaining phenotype, the *rlpA*::stop and Δ*mltC* single mutants exhibited no morphological differences from the WT in either LB or minimal medium (**Fig. S2B**). Therefore, MltC and RlpA collectively fulfill a crucial role in daughter cell separation. Despite visible chaining, the Δ2 LTG mutant grew as well as the WT in batch culture (**Fig. S2C**); however, we did observe a strong motility defect that appeared to be largely due to RlpA inactivation (**Fig. S2D**).

**Fig 2.**
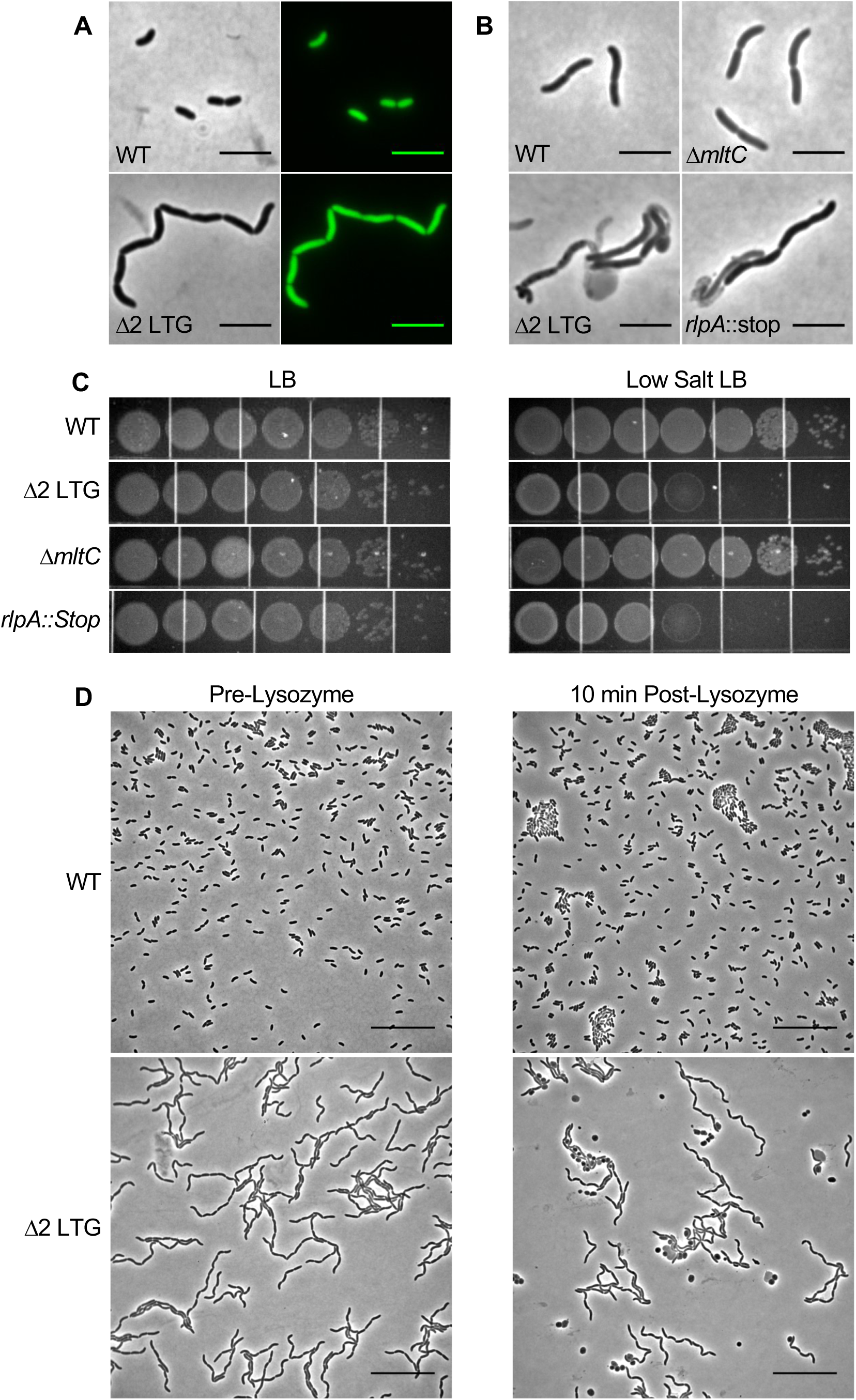
A mutant defective in *rlpA* and *mltC* exhibits a chaining defect. **A)** WT and Δ2 LTG (Δ*mltC rlpA::*stop) cultures were grown to OD_600_ ∼0.8 in LB at 37°C and imaged on agarose pads. Cytoplasmic GFP was expressed constitutively from the native *lacZ* locus. **B)** The indicated strains were grown for 4 hours in low salt LB (0 mM NaCl) at 37°C and imaged on agarose pads. **C)** Overnight cultures of the indicated strains grown in LB were diluted in LB and spot-plated on LB and low salt LB and incubated for 18hrs at 30°C. Grid lines = 1cm. **D)** Exponential phase cultures of WT and Δ2 LTG were exposed to 5 mg/mL lysozyme for 10 min. Scale bars = 25 µm. All data are representative of at least two biological replicates.

It has been previously reported that a mutant *Pseudomonas aeruginosa* lacking RlpA forms unseparated chains of cells when grown in media with low osmolarity (Jorgenson et al., 2014). Similarly, the *V. cholerae rlpA::stop* single mutant, but not the Δ*mltC* single mutant, formed chains of cells when grown in low-salt LB medium and exhibited a 1000-fold plating defect on low-salt LB plates (**Fig. 2BC**). Interestingly, none of these phenotypes were replicated in *Escherichia coli*. An *E. coli* Δ*rlpA* mutant exhibits no obvious morphological defects in LB or low-salt medium (Jorgenson et al., 2014), though this is unsurprising as *E. coli* RlpA, despite being well-conserved, is predicted to be non-functional as a LTG (Jorgenson, 2014). Informed by our *V. cholerae* Δ2 LTG mutant and the *S. enterica* Δ*mltC* Δ*mltE* mutant phenotypes (Wall et al., 2011), we made additional, simultaneous mutations in *E. coli mltC* and *mltE*, but still observed only a very mild morphological defect **(Fig S3)**. Thus, there are species-specific differences in the utilization of these autolysin homologs.

### Δ2 LTG mutations result in outer membrane permeability perturbations

Given that daughter cell separation-defective chains are sensitive to osmotic stress, we also hypothesized that the outer membrane (OM) may be compromised in the Δ2 LTG mutant, as the OM is an important structural and load-bearing component of Gram-negative bacteria (Rojas et al., 2018). Lysozyme typically cannot permeate the OM in sufficient quantities to cause damage to the Gram-negative cell wall. Consistent with this, WT *V. cholerae* tolerated exposure to 5 mg mL^-1^ lysozyme without exhibiting signs of morphological defects (**Fig. 2D**). In contrast, a significant portion of Δ2 LTG mutant cells exhibited a loss of rod shape after just a brief exposure to the same concentration of lysozyme (**Fig. 2D**). This suggests that delayed cell separation may present a barrier to proper OM invagination and results in OM permeabilization.

### Δ2 LTG and Δ*amiB* chaining phenotypes are growth phase dependent

The Δ2 LTG chaining phenotype is reminiscent of a mutant deleted in *V. cholerae’*s sole amidase, AmiB, which also exhibits chain formation (Möll et al., 2014) due to a well-understood daughter cell separation defect (Heidrich et al., 2001, 2002b; Priyadarshini et al., 2007; Uehara et al., 2009, 2010). We thus explored other similarities between the two mutants. During our imaging experiments, we noticed that overnight cultures of Δ2 LTG and Δ*amiB* mutants were devoid of any cell chains, suggesting that chaining was a growth phase-specific phenotype. Single cells of stationary phase Δ2 LTG and Δ*amiB* were not spontaneous suppressor mutants arising at a high frequency, as redilution into exponential phase always resulted in renewed chain formation and subsequent resolution in stationary phase (**Fig. S4**). To more precisely define the growth stage that promotes cell chain resolution in these mutants, we imaged the mutant strains versus WT over their growth cycle and quantified the cells per chain at each time point (**Fig. 3A**). Both Δ2 LTG and Δ*amiB* exhibited peak chain lengths in mid-to late exponential phase (OD_600_ ∼0.8, median chain length of 8 cells for Δ2 LTG and 24 cells for Δ*amiB*), followed by a gradual decline in chain length as the cells entered stationary phase.

**Fig 3.**
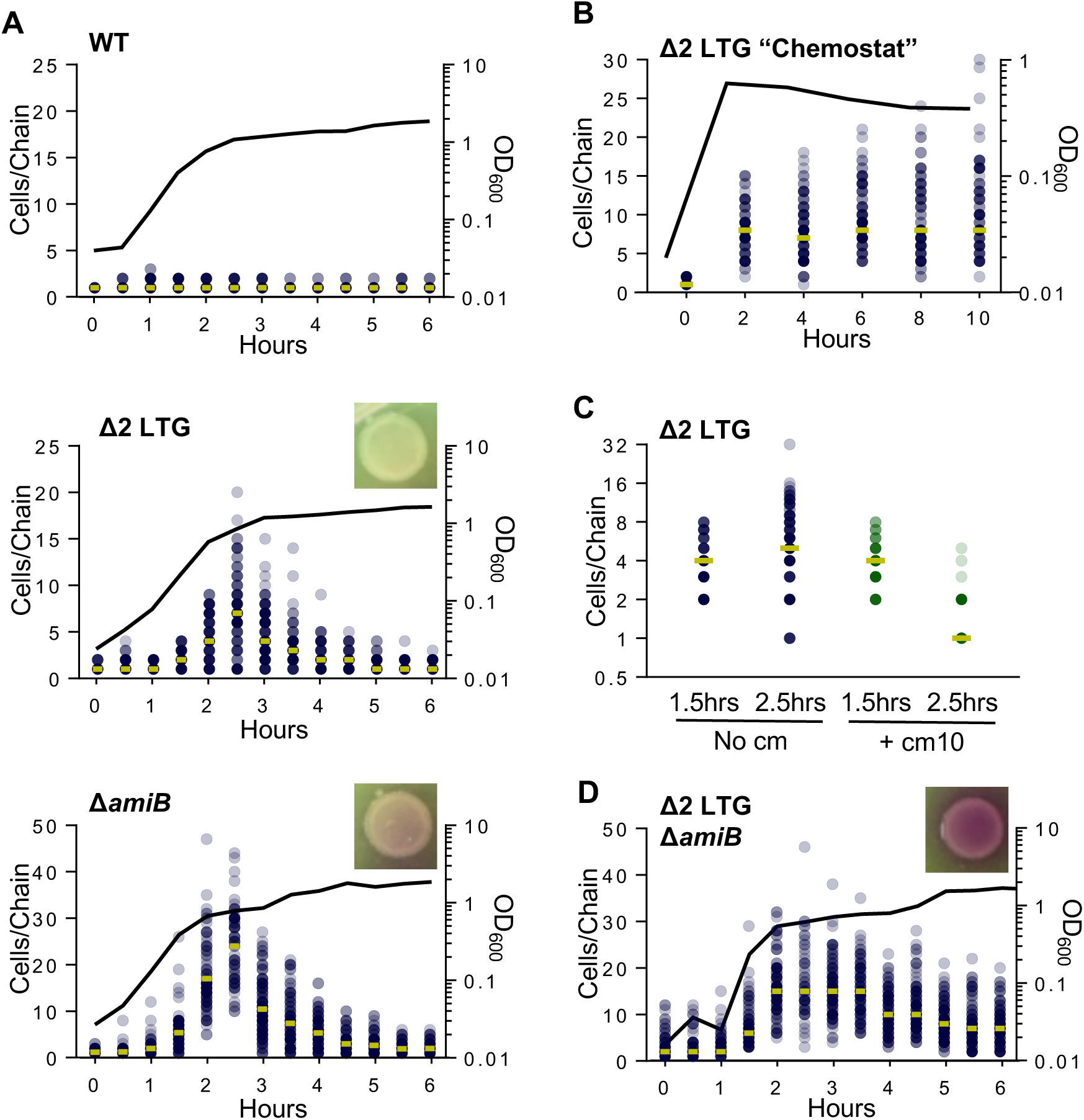
Chaining defects of septal autolysin mutants depend on fast growth and stochastic resolution. **A)** WT, Δ2 LTG (Δ*mltC rlpA::*stop), and Δ*amiB* were grown in LB at 37°C and imaged on agarose pads. Cells per chain where manually counted (n >100). Circles represent raw data points of cells/chain (gold bar = median), line graph shows OD_600_. **B)** A Δ2 LTG culture was back-diluted every 2 hrs into pre-warmed LB at 37°C to maintain exponential phase and imaged on agarose pads. Analysis was conducted as described for Fig. 3A **D)** Δ2 LTG Δ*rpoS* was grown in LB at 37°C and imaged on agar pads. Analysis was conducted as described for Fig. 3A **E)** Chloramphenicol (10 µg/mL, ∼10 × MIC) was added to cultures after growth to OD_600_ ∼0.45 in LB at 37°C and imaged on agarose pads after 1 hr. Analysis was conducted as described for Fig. 3A. **D)** Δ2 LTG Δ*amiB* was grown in LB at 37°C and imaged on agarose pads. Analysis of chain length and CPRG lysis assays were performed as described for Fig. 3A. All data are representative of at least two biological replicates.

The recurring shift between an enrichment of chains in exponential phase and single cells in stationary phase could be explained by stationary phase-specific events, e.g., growth-phase specific lysis cycles or the induction of stationary phase-specific PG remodeling pathways. Alternatively, cell separation could be due to a stochastic process, i.e., reduced septal cleavage activity is initially outpaced by septal PG synthesis during rapid growth, but becomes sufficient for daughter cell separation as division rates slow in stationary phase. Growth phase-dependent chaining has been previously reported for *E. coli* amidase mutants and it was postulated that single cells were likely generated by lysis of cells within a chain (Heidrich et al., 2001). We tested this hypothesis in *V. cholerae* by plating autolysin mutants on CPRG. The Δ2 LTG mutant failed to indicate significant CPRG degradation (**Fig. 3A**), suggesting that lytic elimination of long chains, either by mechanical sheering or cell death, is an unlikely explanation for stationary phase cell separation. The absence of excessive cell debris in microscope images corroborated this observation, as did the WT-equivalent growth rate of this mutant (**Fig. S2C**). In contrast, the Δ*amiB* mutant did exhibit a slight color change on CPRG, indicative of higher background lysis, which is consistent with previous observations in *E. coli* amidase mutants (Heidrich et al., 2001; Priyadarshini et al., 2007). Additionally, this mutant in *V. cholerae* suffers a slight defect in growth rate when compared to WT (Möll et al., 2014) and Δ2 LTG, suggesting that at least some of the chain resolution of the Δ*amiB* mutant may depend on lysis (**Fig. 3A**, **Fig. S2C**). Thus, while cell lysis may somewhat contribute to apparent chain “resolution” in the amidase mutant, it is unlikely to facilitate the same in the Δ2 LTG mutant.

### Δ2 LTG chain resolution is stochastic

We conducted several experiments to help distinguish between chain resolution by stationary phase-induced PG remodeling and stochastic activity of other hydrolases. First, we performed a chemostat-like experiment where Δ2 LTG cells were kept in prolonged exponential phase through periodic back-dilution to maintain an OD_600_ < 0.6. In a simple scenario, if chain resolution were mediated by a stationary phase-exclusive factor, chains would be expected to elongate *ad infinitum* when kept in perpetual exponential phase and single cells would no longer be present in the population. However, if other autolysins could stochastically resolve shared septal PG, albeit with a lower efficiency, we would expect an increase in the variation of chain lengths rather than infinite chains. What we observed was more indicative of stochastic resolution. There was some increase in maximum chain length (**Fig. 3B**), but chains must also have achieved some degree of resolution, as short chains and single cells were still present even after five back-dilutions. These results tentatively suggest that an equilibrium between division and subsequent separation can be achieved in a Δ2 LTG mutant without entering stationary phase.

To test this further, we also surveyed other possible stationary phase-specific factors that could affect cell separation. Culturing Δ2 LTG in the filter-sterilized supernatant of a saturated WT culture failed to prevent chaining (**Fig. S5A**) as would have been expected should chain resolution be modulated by a secreted compound, for example D-amino acids, which in *V. cholerae* are exclusively produced in stationary phase (Cava, De Pedro, et al., 2011; Cava, Lam, et al., 2011). We also inactivated the global stationary phase transcriptional regulator, RpoS, in the Δ2 LTG background and found that chain formation during exponential phase and chain resolution during stationary phase were unaffected (**Fig. S5B**). Finally, we blocked all new protein synthesis in Δ2 LTG grown to exponential phase (OD_600_ ∼0.4) with chloramphenicol (10µg mL^-1^, ∼15 × minimum inhibitory concentration [MIC]). The majority of chloramphenicol-treated Δ2 LTG chains resolved to single or double cells within an hour of chloramphenicol treatment while untreated chains elongated in this time (**Fig. 3C**). This indicates that the cells have already translated the necessary enzymes for chain resolution by OD_600_ 0.4 and adds further support that chain resolution in Δ2 LTG can be achieved by reducing the division rate.

These experiments collectively indicate that the enzymes responsible for septal resolution in the absence of MltC and RlpA are expressed in exponential phase and do not depend on stationary phase-specific signals or factors. Therefore, it is likely that redundant housekeeping PG hydrolases or LTGs can mediate daughter cell separation, albeit at a lower efficiency, when the main separation systems are inactivated. We thus hypothesize that chain resolution during transition into stationary phase is the consequence of reduced division rate at this growth stage, which allows other, less efficient, autolysins to “catch up” and separate daughter cells.

### Alternative septal resolution factors are insufficient in the cumulative absence of RlpA, MltC, and AmiB

Given the similarities between the Δ2 LTG and Δ*amiB* mutants and the established dependence of cell separation on the tight spatiotemporal regulation of AmiB (Priyadarshini et al., 2006, 2007; Uehara et al., 2010; D. C. Yang et al., 2011; Peters et al., 2011; Möll et al., 2014), we tested the formal hypothesis that RlpA and MltC directly or indirectly contributed to AmiB recruitment and that the Δ2 LTG chaining defect might thus be due to lack of AmiB localization. A functional AmiB-mCherry fusion localized to septal rings in Δ2 LTG chains, suggesting that this cell separation defect occurs despite proper AmiB localization (**Fig. S6AB**).

We then investigated the possibly redundant autolytic roles of AmiB, RlpA, and MltC by generating a Δ2 LTG Δ*amiB* triple mutant and quantifying growth phase-dependent chaining. While this strain was viable, it grew much more slowly than the Δ2 LTG or Δ*amiB* mutants (**Fig. S2C**) and perhaps more strikingly, chain resolution was incomplete even after 24 hours and gave visual evidence of strong lysis under the microscope as well as on CPRG plates (**Fig. 3D**, **Fig. S4**). This increased lysis and failure to completely resolve chains suggest that the amidase and septal LTGs are the principle daughter cell separation systems.

We were interested to learn whether the additive, deleterious effects of the Δ2 LTG and Δ*amiB* mutations were due to the unique potential functions of LTGs versus amidases at the septum, or if over-expression of AmiB, or other classes autolysins for that matter, could complement the Δ2 LTG chaining defect. Overexpression of *V. cholerae*’s primary housekeeping endopeptidases, ShyA or ShyC (Dörr et al., 2013), was unable to appreciably reduce chaining in the Δ2 LTG mutant (**Fig. 4A**) (functional expression of these constructs was validated by their ability to complement a mutant defective in multiple endopeptidases (Dörr, Davis, et al., 2015)). Similarly, overexpression of *V. cholerae*’s only amidase, AmiB (co-overexpressed with its activators NlpD and EnvC [Möll et al., 2014; Uehara et al., 2009; Yang et al., 2011]), demonstrated functionality by complementing the Δ*amiB* mutant (**FigS6B**), but could not prevent chaining in the Δ2 LTG mutant. Thus, daughter cell separation likely specifically requires LTG activity, rather than increased general PG hydrolysis.

**Fig 4.**
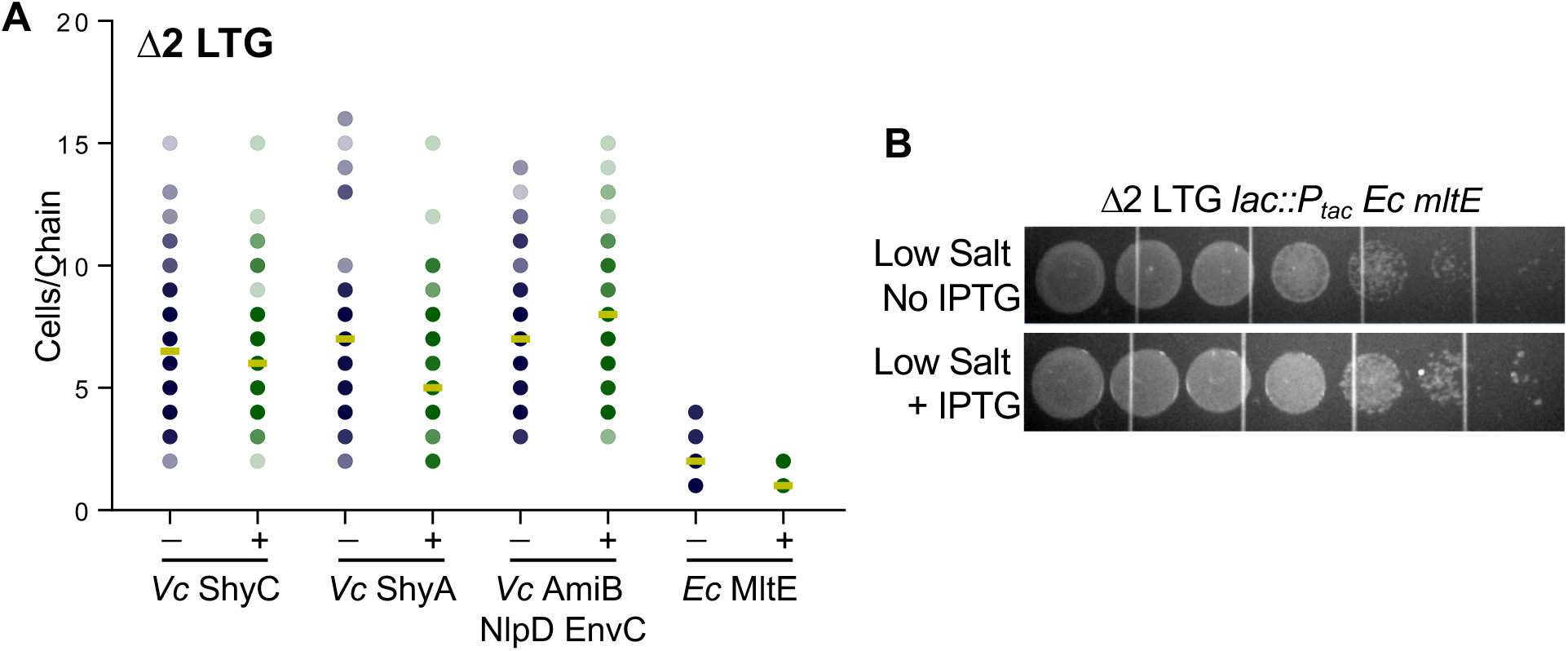
Lytic transglycosylase activity is required for septal PG resolution. **(A)**Expression of chromosomal P_tac_ : *shyC*, P_tac_ : *shyA*, P_tac_ : *amiB nlpD envC, or* P_tac_ : *mltE*_*E. coli*_ was induced with 1 mM IPTG in a Δ2 LTG background, grown in LB at 37°C to ∼OD_600_ 0.6, and imaged on agarose pads. Analysis was conducted as described for Fig. 3A **(B)**An overnight culture of Δ2 LTG P_tac_ : *mltE*_*E. coli*_ grown in LB was diluted in LB and spot-plated on low salt LB +/- 1mM IPTG and incubated for 18hrs at 30°C. Grid lines = 1cm. All data are representative of at least two biological replicates.

Interestingly, we found that heterologous expression of the *E. coli* LTG MltE was highly effective at facilitating septal resolution in the Δ2 LTG mutant (**Fig. 4A**). MltE has a relatively broad spectrum of PG substrate specificity, shown *in vitro* to generate products indicative of both endo-and exolytic cleavage on denuded or un-crosslinked muropeptides (M. Lee et al., 2013; Dik et al., 2017; Byun et al., 2018), and has no known homolog in *V. cholerae.* Overexpression of *mltE* was not toxic in a WT background, suggesting that it primarily digests septal PG in the *V. cholerae* Δ2 LTG mutant (**Fig. S7**). MltE has a strong preference for uncrosslinked PG (M. Lee et al., 2013; Dik et al., 2017; Byun et al., 2018), providing a possible explanation for its lack of general toxicity, since the main body of the cell’s PG is generally crosslinked (Desmarais et al., 2015). Incidentally, this also suggests that septal PG in Δ2 LTG is largely uncrosslinked; which would be consistent with completed or concurrent amidase or endopeptidase activity. MltE was also able to rescue Δ2 LTG growth in low-salt media (**Fig. 4B**). In combination, these observations suggest that some characteristic of the division septum requires the activity of LTGs over other autolysins to allow for septal resolution and daughter cell separation.

### RlpA and MltC are late division proteins

Division proteins are often recruited specifically to midcell. We generated stable, functional **(Fig. S8AB)** translational fusions of MltC and RlpA to mCherry to track localization throughout growth and division. Both RlpA-mCherry and MltC-mCherry clearly localized to the midcell and co-expression of FtsZ-YFP or YFP-FtsN further revealed that both LTGs arrive closer to or after FtsN recruitment to the septum, indicating that RlpA and MltC are likely late division proteins (**Fig. 5**). RlpA midcell localization is consistent with the published roles of its highly conserved homologues in *E. coli* and *P. aeruginosa* (Gerding et al., 2009; Jorgenson et al., 2014; Yahashiri et al., 2015), however, the subcellular localization pattern of MltC homologs in any organism has not been previously reported.

**Fig 5.**
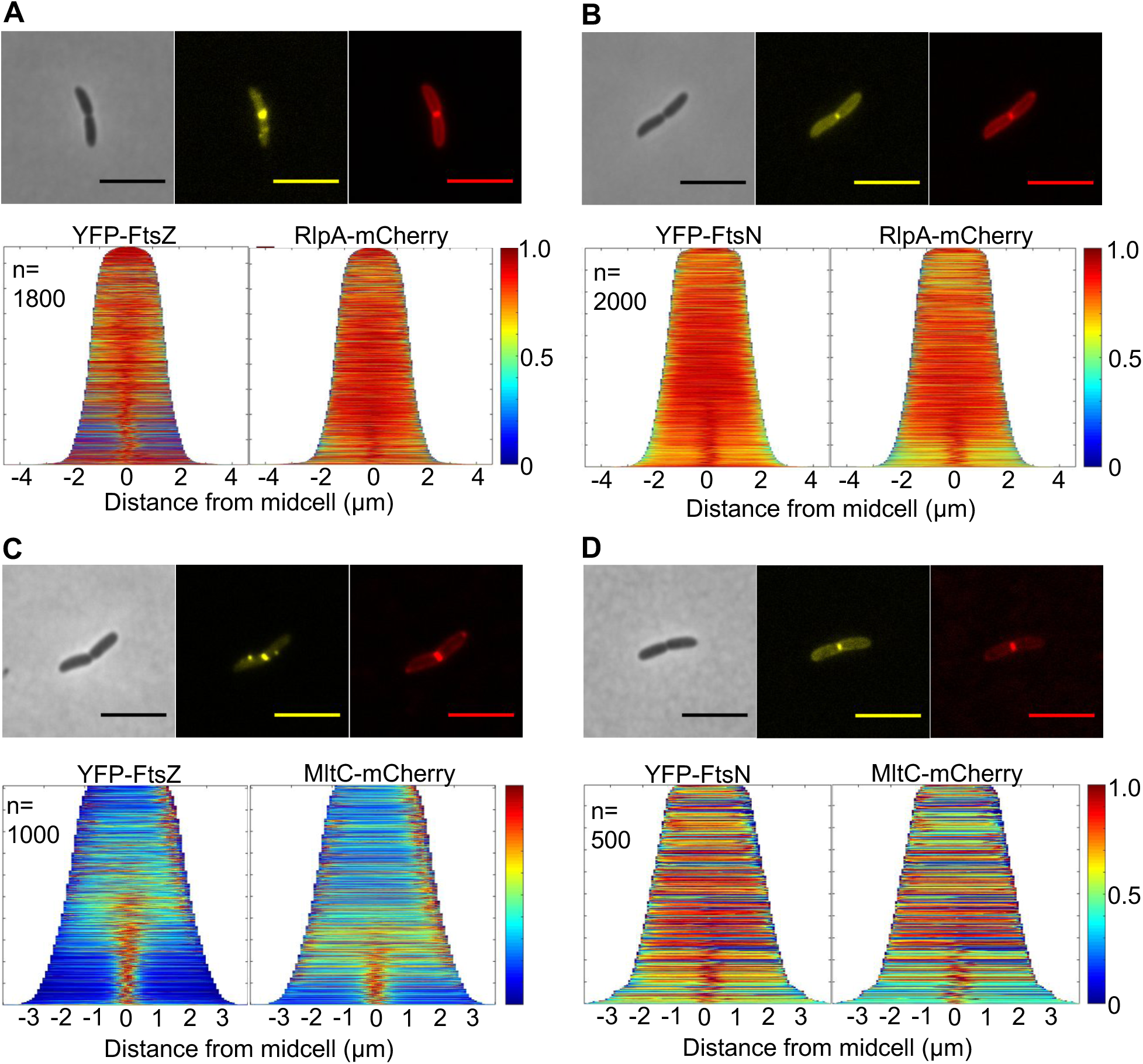
RlpA and MltC are recruited to the septum during late stages of division. WT carrying pBAD33 *yfp-ftsN* or *yfp-ftsZ* and pHL100 *rlpA-mCherry* or *mltC-mCherry* was grown in M9 + 0.2% glucose supplemented with the appropriate antibiotics at 30°C, induced with 0.2% arabinose and 1mM IPTG after 2hrs, and imaged on agarose pads at OD_600_ ∼0.15. Demographs of **A)** RlpA/FtsZ co-localization, **B)** RlpA/FtsN co-localization,**C)** MltC/FtsZ co-localization, and **D)** MltC/FtsN co-localization were generated using Oufti. All data are representative of at least two biological replicates.

### Outer membrane insertion is not required for MltC and RlpA LTG activity

Both LTGs are predicted outer membrane proteins with well-conserved outer membrane target lipoboxes LXGC (**Fig. S9A)** (Babu et al., 2006), so we postulated that this localization may be important for either their recruitment to the midcell or for their function as septal cleavage enzymes. To test this, we substituted the outer membrane targeting signal sequences of MltC and RlpA N-terminal mCherry fusions with the periplasmic signal sequence of thiol disulfide oxidoreductase, DsbA. Surprisingly, these soluble versions of both proteins were able to complement the Δ2 LTG chaining phenotype in LB (**Fig. 6A**) yet did so with what appeared to be reduced localization to the midcell (**Fig. S9B**). Thus, outer membrane attachment is not essential to the function of RlpA and MltC in LB. However, the functionality of DsbA^ss^-RlpA_[18-263]_ was not absolute under all conditions; expression of this construct in a Δ2 LTG background during growth in low salt LB resulted in the formation of severe morphological aberrations, including short filaments and bulging at the midcell (**Fig. 6B**). Induction of this defect was not dominant as the DsbA^ss^-RlpA_[18-263]_ mutant did not affect the morphology of WT in low salt LB, nor did over-expression of RlpA-mCherry retaining its native signal sequence affect the morphology of the Δ2 LTG mutant in low salt LB (**Fig. S9CD**). Despite the apparent division defect induced by soluble DsbA^ss^-RlpA_[18-263]_ expression, the mis-localized protein could still restore growth of the Δ2 LTG mutant on low salt LB (**Fig. 6C**). The phenotype induced by the DsbA^ss^-RlpA_[18-263]_ mutant in low-salt medium was more similar to filamentous division mutants than to other chaining autolysin mutants. Though we did not explore this further, we suspect that RlpA may have conserved roles in division other than septal resolution, roles that may explain why *E. coli* retains an LTG-deficient homologue of RlpA.

**Fig 6.**
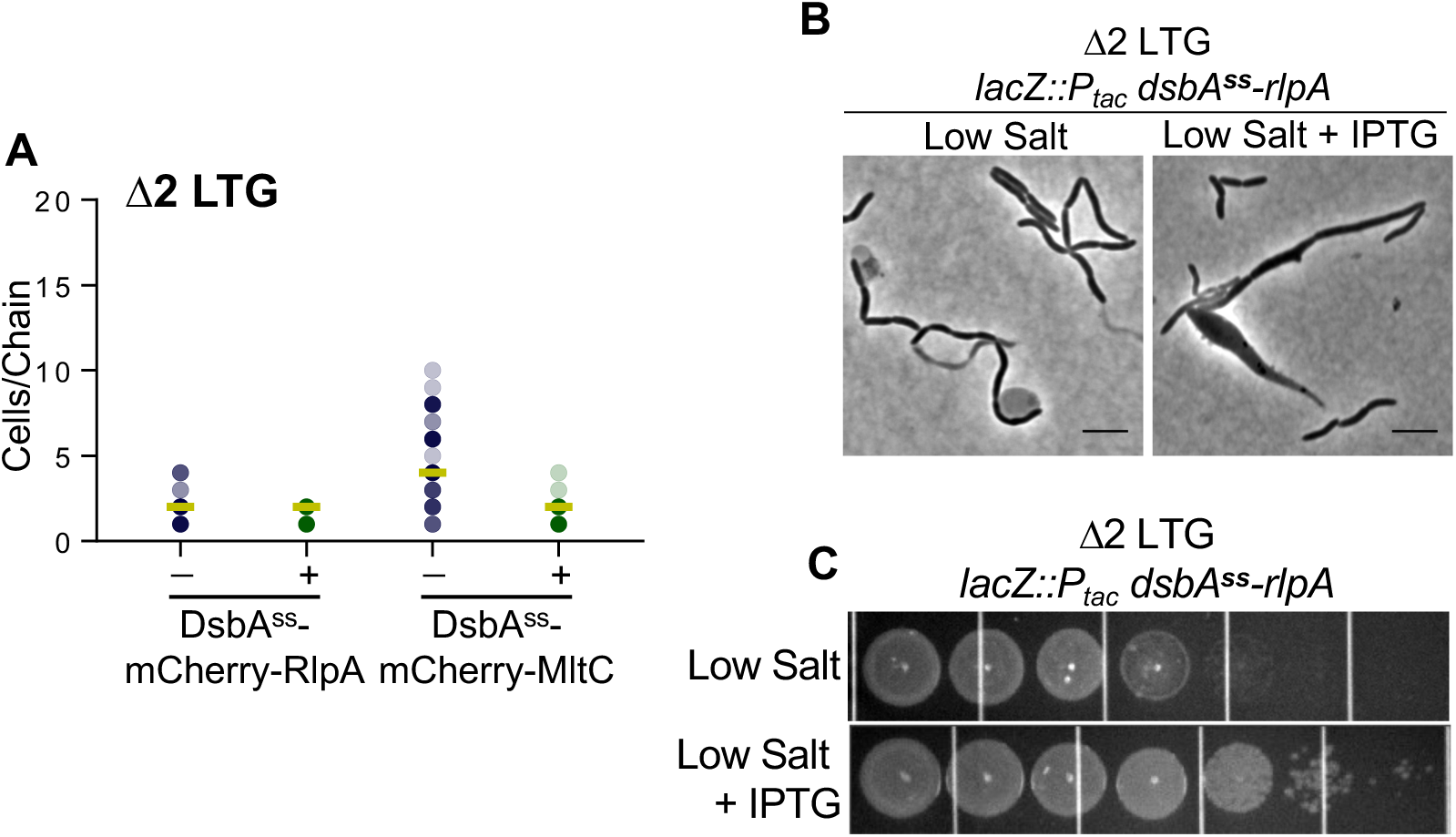
OM insertion of RlpA and MltC is not required for septal resolution. **A)** Expression of chromosomal P_tac_ : *dsbA*^*ss*^*-rlpA*_[18-263]_ or *dsbA*^*ss*^-*mltC*_[42-396]_ was induced with 1 mM IPTG in a Δ2 LTG background, grown in LB at 37°C to ∼OD_600_ 0.6, and imaged on agarose pads. Analysis was conducted as described for Fig. 3A **B)** Δ2 LTG P_tac_ :*dsbA*^*ss*^*-rlpA*_[18-263]_ was grown in low salt LB +/- 1 mM IPTG at 37°C for 4 hrs and imaged on agarose pads. **C)** Overnight culture of Δ2 LTG P_tac_ : *dsbA*^*ss*^*-rlpA*_[18-263]_ grown in LB at 37°C was spot-plated onto low salt LB +/- 1mM IPTG and incubated overnight at 30°C. Grid lines = 1cm. All data are representative of at least two biological replicates.

## DISCUSSION

### LTG activity plays a conserved role in septal PG resolution

We report here that two LTGs, RlpA and MltC, are septum-specific enzymes with crucial roles in daughter cell separation in *V. cholerae.* To our knowledge, this is the first report of a physiological role for any housekeeping LTG in the cholera pathogen and the second direct report of a physiological role for MltC (Wall et al., 2011). Follow-up studies will be required to determine how septal LTGs interact with the divisome and the PG structural reasons for why MltC and RlpA specifically contribute so significantly to the separation of daughter cells. RlpA and MltC are well-conserved amongst proteobacteria (Artola-Recolons et al., 2014; Yahashiri et al., 2015) yet despite this conservation, mutations in either RlpA (Jorgenson et al., 2014), MltC, or both fail to produce phenotypes in the classic model system *E. coli.* This study serves to highlight the importance of variety in microbial perspective to understand key processes in bacterial physiology. At the same time, the high conservation of RlpA, especially amongst Gram-negatives (Yunck et al., 2016), and the aberrant morphology of a Δ2 LTG mutant overexpressing *dsbA*^*ss*^*-rlpA* in low salt suggests that septal LTGs may also have other unappreciated roles in bacterial cell division and daughter cell separation in other Gram-negative pathogens.

### A “Cleave and Clear” LTG-dependent model of daughter cell separation

With the addition of RlpA and MltC to the division complex, we can add more detail to the molecular landscape of the septal cell wall. We observed that the chaining defect of the Δ*amiB* mutant was slightly more severe than the Δ2 LTG mutant, suggesting that the majority of PG shared between daughter cells at the septum is connected by peptide crosslinks that can be removed by amidase activity. However, the chaining phenotype of the Δ2 LTG mutant implies that there is shared septal PG that cannot be resolved by amidase activity. It is ideally expected, from the close association of septal PG synthesis with rotating FtsZ filaments, that PG strands are inserted perpendicular to the long cell axis (Daniel & Errington, 2003; X. Yang et al., 2017; Bisson-Filho et al., 2018). Our findings lead us to speculate that the cell division machinery stochastically generates PG strands that are deposited and/or elongated at a non-ideal angle such that they transect the septal plane to create a bridge between daughter cells (**Fig. 7**). Since amidase activity is precisely controlled by activators NlpD and EnvC (Möll et al., 2014), the radius of amidase activity is likely highly restricted. Some aberrantly deposited glycan strands might then be crosslinked outside of the range of amidase activity, requiring LTGs as backup enzymes to cleave connecting PG left behind after amidase activity is complete. Consistent with this idea, RlpA in *P. aeruginosa* acts endolytically on denuded glycan strands (Jorgenson et al., 2014; M. Lee et al., 2017) and may follow AmiB to cleave remaining bridging strands. MltC may function to resolve septal PG in two ways. It might also perform a similar function to RlpA, as some endolytic activity has been attributed *E. coli* MltC (Artola-Recolons et al., 2014). One might also envision that long PG strands between daughter cells, even those that are no longer covalently linked to the PG network, might still present an obstacle to outer-membrane invagination. MltC, which in *E. coli* has been shown to also be processive and exolytic, but inactive on crosslinked PG strands, could be responsible for clearing such debris from the path of outer-membrane invagination (Artola-Recolons et al., 2014).

**Fig 7.**
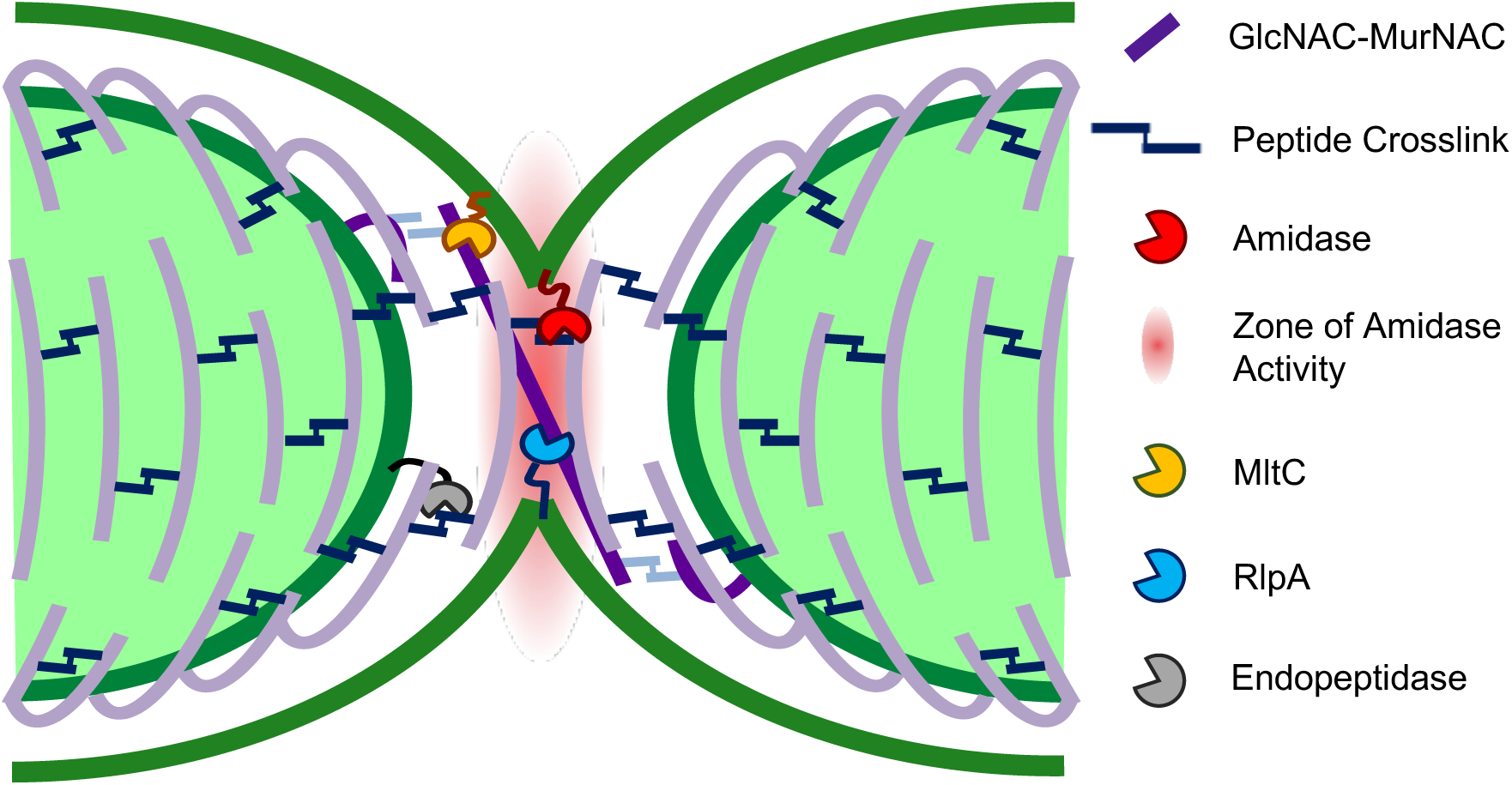
Model of RlpA and MltC requirement for daughter cell separation. Septal LTGs may remove PG debris that cannot be processed by other autolysins and that impedes the completion of OM invagination.

We suggest that a function to remove aberrant septal PG strands would be particularly important for *V. cholerae*, which completes the final stages of cell division with astonishing speed (Galli et al., 2017). While the model bacteria *E. coli, B. subtilis*, and *C. crescentus* have mature divisomes by 50% of the cell cycle (Aarsman et al., 2005; Gamba et al., 2009; Goley et al., 2011) and *E. coli and B. subtilis* both encode multiple amidases to assist in daughter cell separation (Heidrich et al., 2001; Firczuk & Bochtler, 2007), *V. cholerae* encodes a single amidase and septation occurs only in the final 10% of its cell cycle (Galli et al., 2017) This is likely to increase the chance for errors to occur, which would create a need to efficiently remove spatial obstructions, including PG aberrantly crosslinked or uncrosslinked PG debris. The absence of a strong chaining phenotype in *E. coli* Δ*mltC* Δ*rlpA* Δ*mltE* (and the need to delete further LTGs to elicit a weak version of such a phenotype) may indeed reflects this species’ slower division process, resulting in fewer errors. It should also be noted that repression of P_tac_ is somewhat leaky (Rosano & Ceccarelli, 2014) and that complementation of either the Δ2 LTG or Δ*amiB* chaining defects occurred readily without induction, which suggests that very little autolytic activity is actually required for separation and/or that obstruction of OM invagination by PG is rare.

The exponential phase-dependent manner of the Δ*amiB* and Δ2 LTG chaining phenotype suggests that AmiB and RlpA/MltC functions are redundant with at least one other autolysin whose activity alone is insufficient to sustain daughter cell separation at the fast rate imposed by exponential growth. We hypothesize that RlpA/MltC and AmiB maintain separate functions and that chain resolution in stationary phase is a reflection of each system requiring a distinct autolytic substitute (or group of autolysins) to mediate chain resolution in its absence. For the amidase, this substitute could be D,D or L,D endopeptidases, for LTGs the substitute could be other remaining LTGs. Thus, RlpA, MltC, and AmiB compose the primary group of autolysins responsible for septal resolution and other LTGs or endopeptidases that are secondary to RlpA/MltC or AmiB, respectively, cannot support daughter cell separation in the absence of all three primary septal autolysins. Future work will address the intricate redundancy relationships between these diverse groups of autolysins.

## EXPERIMENTAL PROCEDURES

### Strains, Media, and Growth Conditions

All *V. cholerae* strains in this study are derivatives of *V. cholerae* WT El Tor strain N16961 (Heidelberg et al., 2000) and *E. coli* strains are derivatives of *E. coli* K-12 strain MG1655 (Blattner et al., 1997).

Strains were grown at 30°C or 37°C in Luria-Bertani (LB) broth with or without 1.5% NaCl, or in M9 medium containing 0.2% glucose (“M9 Salts,” 2006; “M9 minimal medium (standard),” 2010). When required for selection or plasmid retention, growth media were supplemented with 5-Bromo-4-Chloro-3-Indolyl β-D-Galactopyranoside (X-gal, 40 µg mL^-1^) streptomycin (200 µg mL^-1^), kanamycin (5 µg mL^-1^), carbenicillin (100 µg mL^-1^), and/or chloramphenicol (5 µg mL^-1^ for *V. cholerae* and 20 µg mL^-1^ for *E. coli*). Sucrose counter-selection was performed on 10% sucrose LB medium without NaCl. Genes under *P*_*tac*_ or *P*_*BAD*_ control were induced with 1 mM isopropyl-β-D-1-thiogalactopyranoside (IPTG) or 0.2% L-arabinose, respectively. For lysis assays, growth media were supplemented with chlorophenol red-β-D-galactopyranoside (CPRG, 20 µg mL^-1^, Sigma Aldrich # 10884308001).

At least two replicates were completed per strain and condition for growth curves and growth dependent chaining experiments. For growth curves, strains were inoculated 1:100 into 200 µL of medium from an overnight culture and grown in a Bioscreen growth plate reader (Growth Curves America) at 37°C with random shaking of medium amplitude, and OD_600_ readings at 5-minute intervals. For chaining experiments with more than three time points, overnight cultures were diluted 1:100 into 200 mL of medium in a 500 mL non-baffled flask and incubated at 37°C with shaking. For experiments with three or fewer time points, overnight cultures were diluted 1:100 into 5 mL of medium in culture tubes.

### Construction of Plasmids and Strains

All strains are derivatives of *V. cholerae* El Tor N16961 or *E. coli* MG1655. Strains, plasmids, and primers are summarized in **Table S1-S4**. *E. coli* DH5*α λ*pir was used for general cloning while *E. coli* SM10 *λ pir* or MFD *λ pir* (Ferrières et al., 2010) were used for conjugation into *V. cholerae*. Plasmids were constructed using Gibson assembly (Gibson et al., 2009).

*V. cholerae* chromosomal in-frame deletions were generated by amplifying 500bp up-and downstream of the gene of interest by PCR, cloning into suicide vector pCVD442 (Donnenberg & Kaper, 1991), and conjugating into *V. cholerae* N16961 followed by sucrose counter-selection. Briefly, conjugation was performed by mixing and pelleting equal volumes LB overnight culture of plasmid donor *E. coli* SM10 *λpir* or MFD *λpir* strains with recipient *V. cholerae*, spotting mixed pellet onto an LB (+ 600µM diaminopimelic acid for MFD *λpir*), and incubating at 37°C for 3 hrs. Single colonies were first selected for on LB + sm200 + vector selection, followed by counter-selection on salt-free LB + 10% sucrose and PCR verification of the deletion.

Ectopic, inducible chromosomal expression of proteins was achieved by cloning the gene(s) of interest open reading frame with either the native 20bp upstream sequence, a strong, consensus ribosome binding site (RBS), or no ribosome binding site into pTD101, a derivative of pJL1 (Miyata et al., 2013) engineered to carry the lac promoter, *lacIq*, and a multiple cloning site for integration of expression constructs at the native *lacZ* locus. In particular, pAW51 for polycistronic co-expression *amiB, nlpD*, and *envC*, was designed such that *amiB* was translated from a strong consensus RBS and the 20bp upstream regions containing the native RBS’s of *nlpD* and *envC* gene were included between *amiB* and *nlpD* or *nlpD* and *envC*, respectively, to generate a 3-gene transcriptional fusion under P_tac_ regulation. pTD101 derivatives were conjugated into *V. cholerae* as described above for pCVD442.

Depletion of RlpA was accomplished by using suicide vector pAM299 (Möll et al., 2015) to place *rlpA* under P_BAD_ control at its native locus. pAM299 was conjugated into *V. cholerae* as a single crossover, without counterselection. For fluorescent localization or detection of proteins by Western blot, genes were cloned into pBAD33 (Guzman et al., 1995) or pHL100 (Dörr, Alvarez, et al., 2015)

*E. coli* chromosomal in-frame deletions were generated using a combination *λ* of Red and FLP recombinase systems as previously described (Cherepanov & Wackernagel, 1995; Wanner, 2000; Murphy & Campellone, 2003). Briefly, plasmids pKD3 and pKD4 were used as templates for amplifying *cmR* and *kanR* cassettes flanked by FLP recombinase sites with 50bp homology to the up-and downstream regions of the gene of interest. Electrocompetent MG1655 carrying the λRed recombinase system on pKM208 was transformed with the PCR product amplified from pKD3/4 and recombinants selected for on LB + kan50 or cm20. The gene replacements were moved into a clean MG1655 background by P1 phage transduction (Thomason et al., 2007) and the resulting strain transformed with pCP20 carrying the FLP recombinase. Finally, pCP20 was induced and cured at 37°C and candidates screened for loss of antibiotic resistance and PCR verification of the chromosomal gene deletion.

### Microscopy

Cells from liquid culture were imaged without fixation on 0.8% agarose pads containing the same medium from the relevant experiment using a Leica DMi8 inverted microscope. Phase contrast images of chaining mutants were analyzed manually to calculate the number of cells per chain in >100 chains. Raw phase and fluorescent images were analyzed in Oufti (Paintdakhi et al., 2016) using the pre-set *E. coli* M9 subpixel parameters for calculation of cell width and length and for generation of fluorescent signal localization “demographs. ”

### Western Blot Analysis

Expression of translational mCherry fusions was induced in WT *V. cholerae* with 0.2% arabinose for pBAD33 or 1 mM IPTG for pHL100 and *lacZ::P*_*tac*_ in LB and grown to OD_600_∼0.6. Cells were harvested by centrifugation (9500 *g*, 15 min) at room temperature and resuspended in 1% SDS + 10 mM DTT lysis buffer. Resuspended cells were incubated at 95°C for 3 min, then sonicated 4 × 5 seconds at 20% amplitude. Standard Western blots against mCherry were performed using polyclonal mCherry antibody (Genetex #GTX59788) and detection by IRDye 800CW secondary antibody (Li-cor #926-32211). After imaging for mCherry, the same blots were then re-incubated with monoclonal RpoA antibody (BioLegend # 663104) detected by IRDye 800CW secondary antibody on an Odyssey CLx imaging device (Li-cor).

## ACKNOWLEDGEMENTS

We thank Matthew Jorgenson (University of Arkansas) and all members of the Dörr lab for helpful discussion. We also thank the Waldor lab for sharing plasmids pBAD33 *yfp-ftsZ* and pBAD33 *yfp-ftsN* as well as the faculty, staff, and students at the Weill Institute for Cell and Molecular Biology (WICMB) for sharing equipment and resources. Research in the Dörr lab is supported by the NIH/NIAID (1R01 AI143704-01).

## AUTHOR CONTRIBUTIONS

AIW and TD have made major contributions to the design of the study, the collection, analysis, and interpretation of data, and writing of the manuscript. VJR, ST, and BR have made major contributions to the collection and analysis of data. KW has made major contributions to the analysis of data.

