## Supplementary figures and images for "Lytic transglycosylases RlpA and MltC assist in *Vibrio cholerae* daughter cell separation"

### Supplemental Figures

**Fig S1**

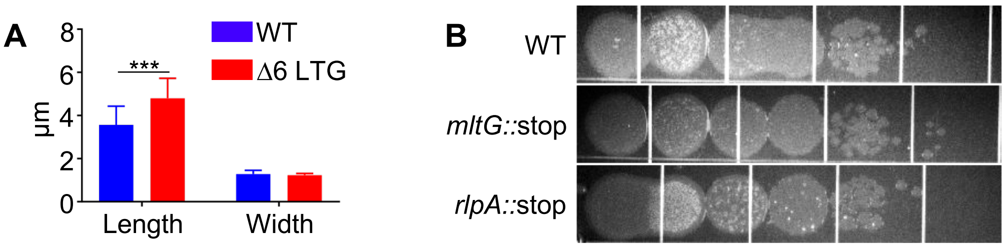

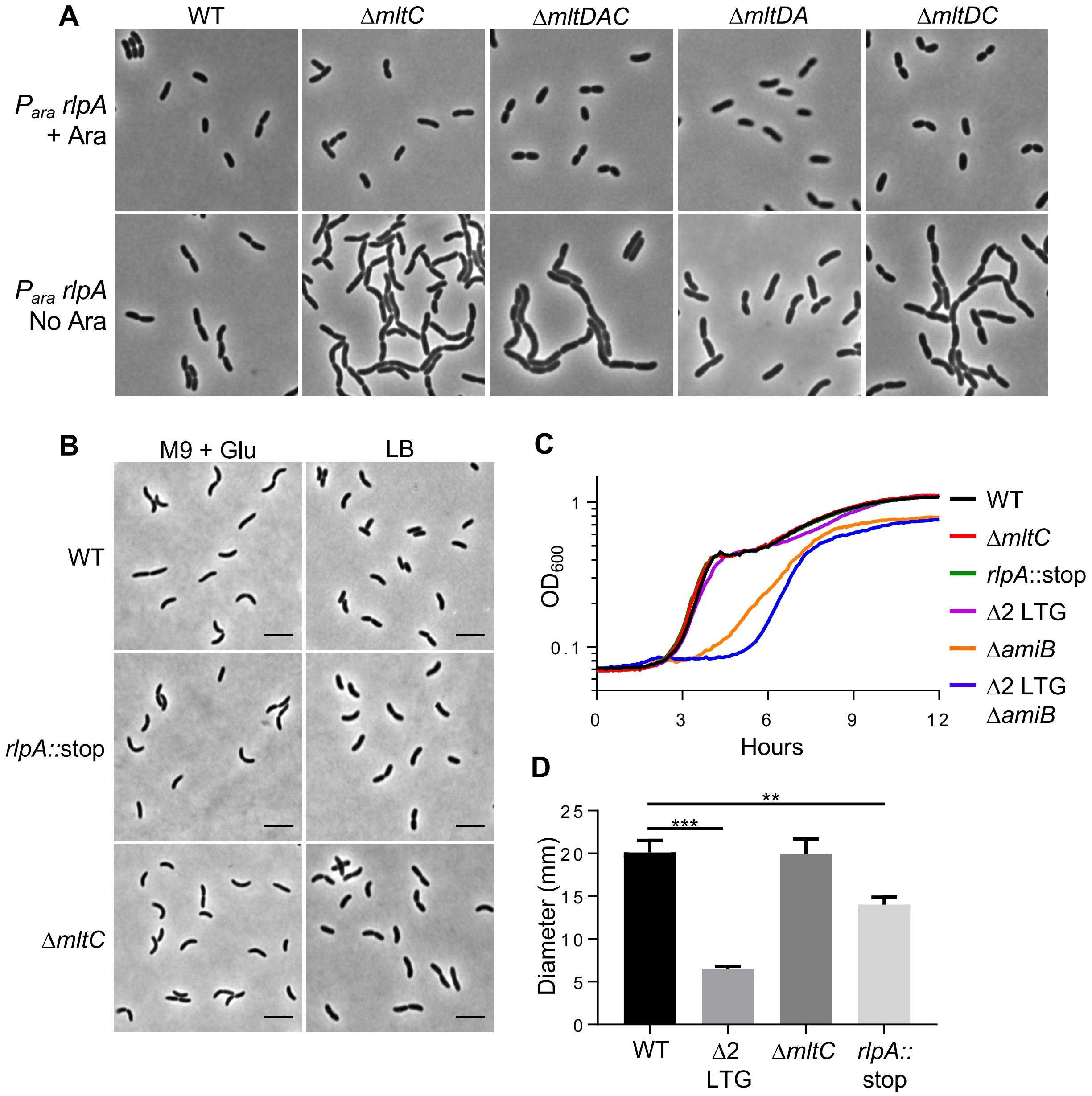

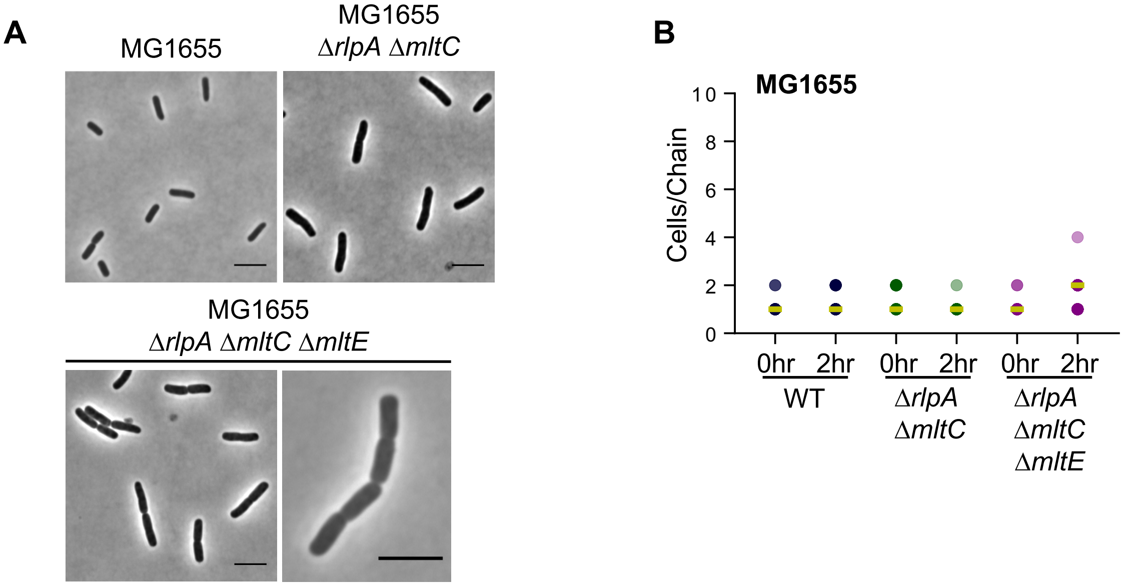

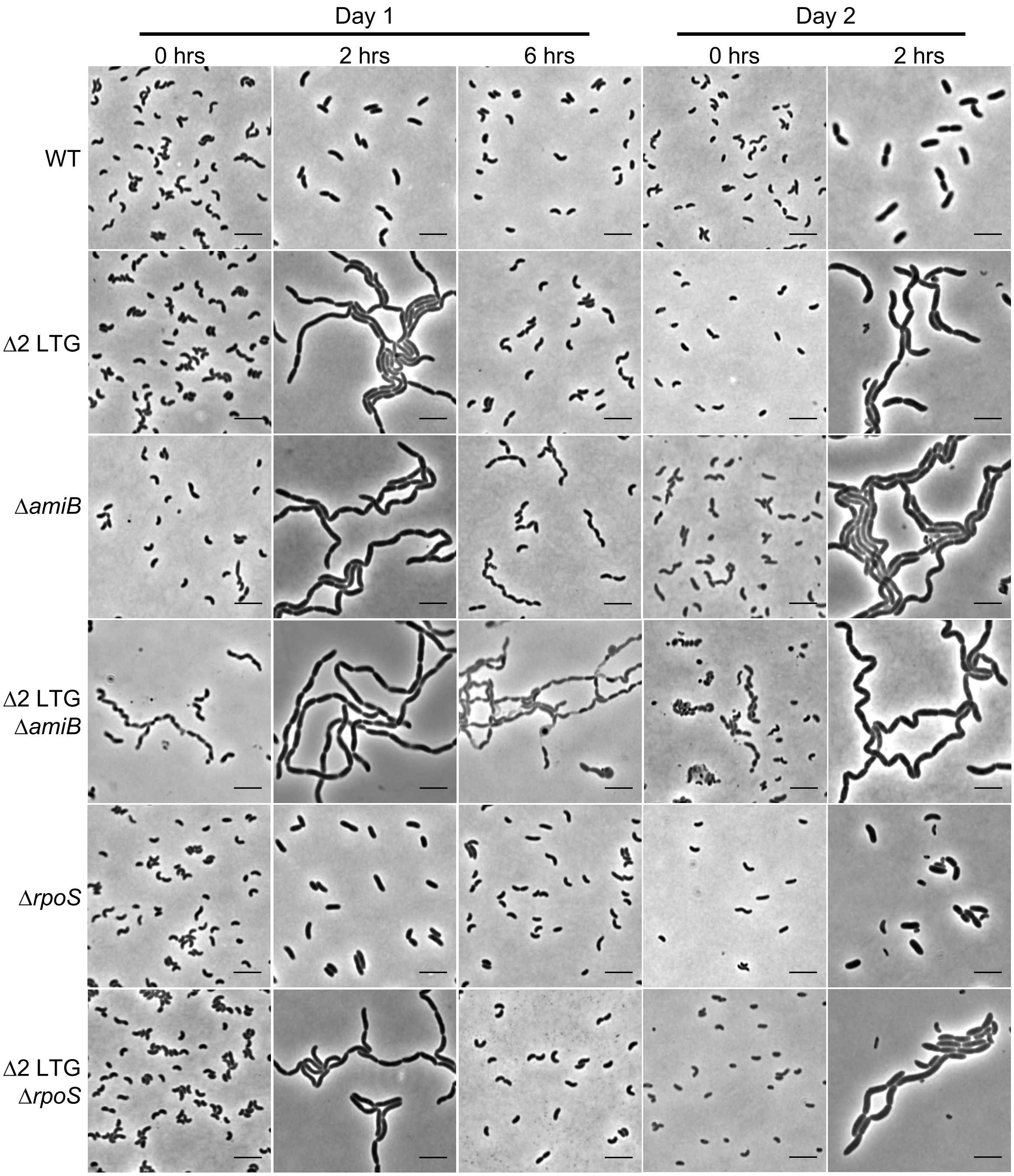

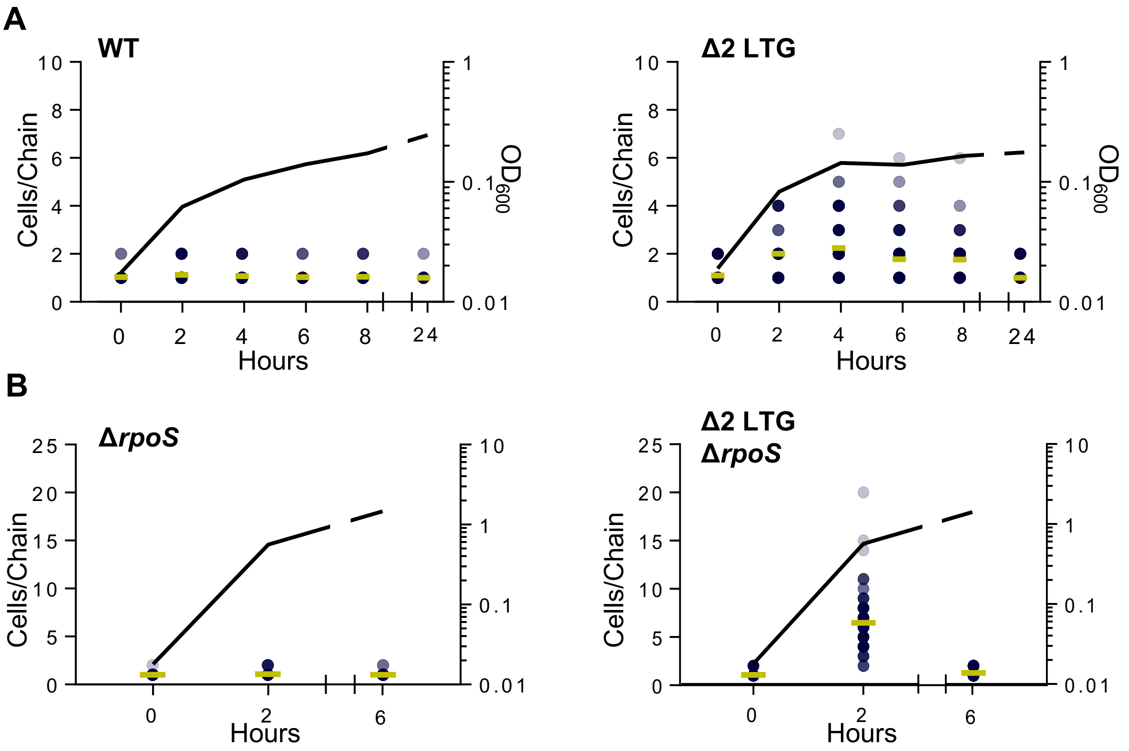

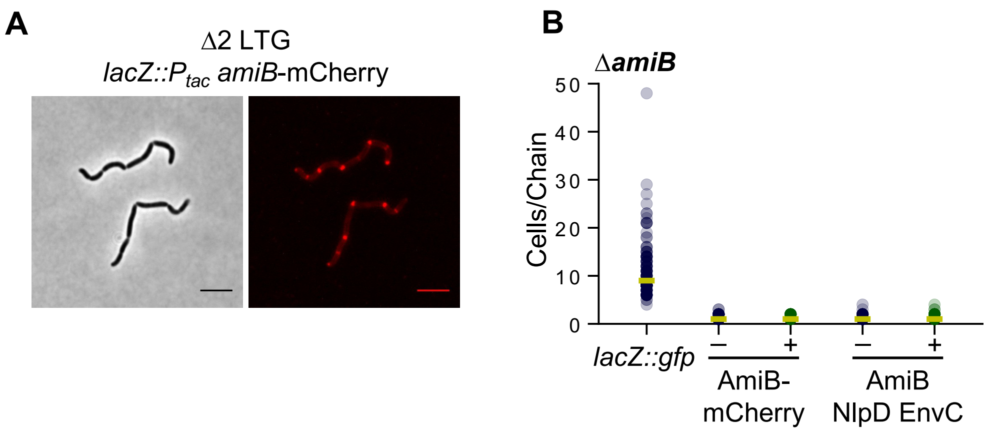

**Fig S7**

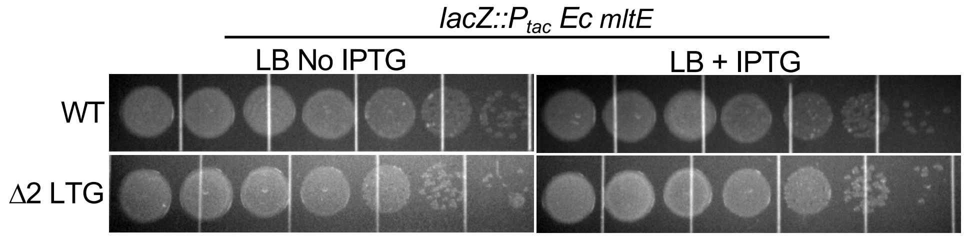

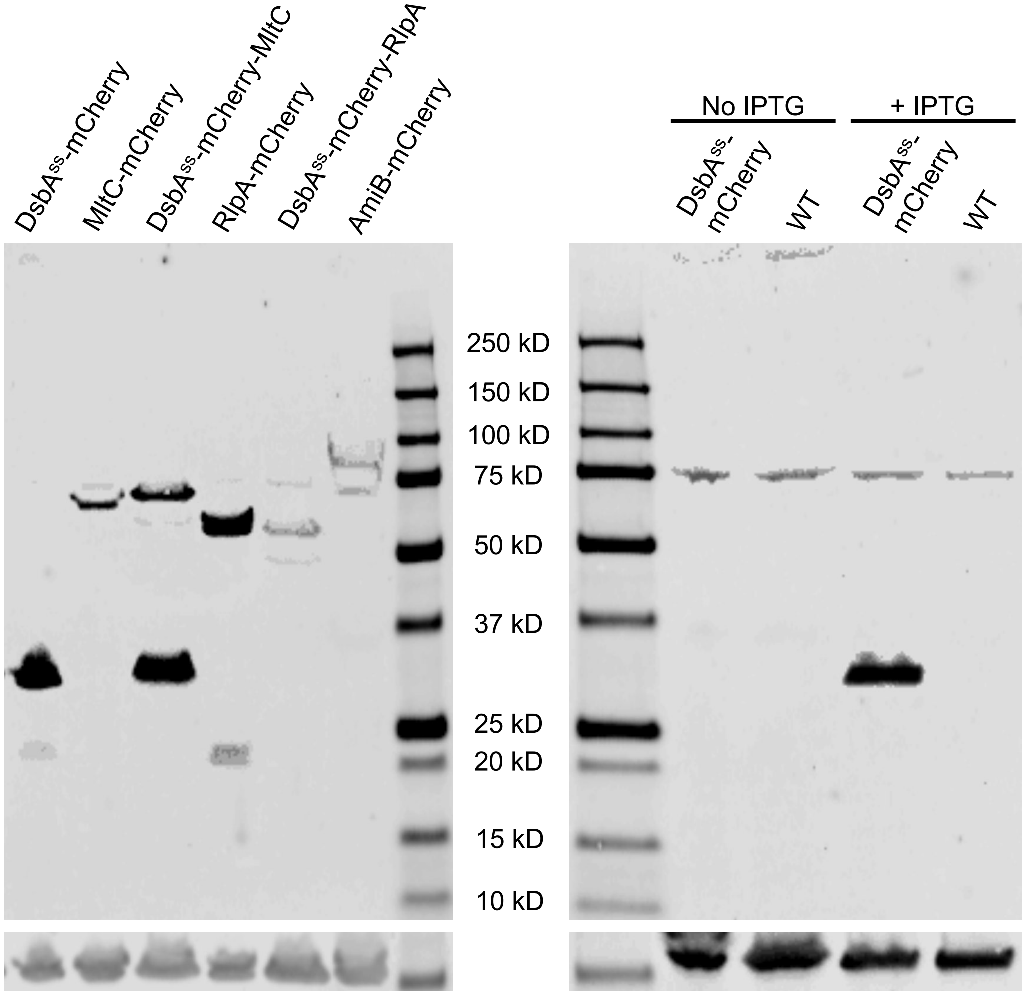

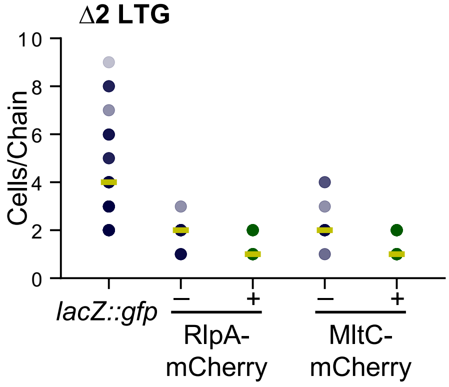

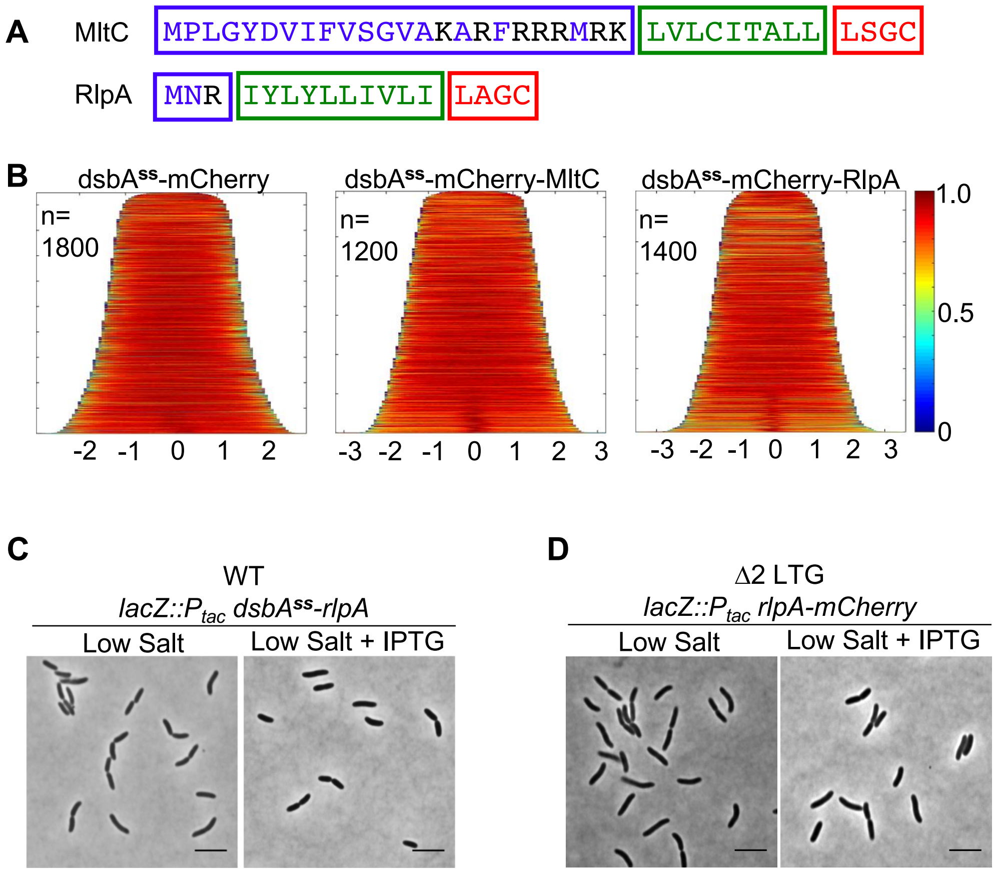
