## Supplemental Tables for "Lytic transglycosylases RlpA and MltC assist in *Vibrio cholerae* daughter cell separation"

**Table S2.** ***E. coli*** **strains**

| Strain | Description | Source or reference |
|---|---|---|
| MG1655 | *E. coli* K-12 strain | Blattner, et. al., 1997 |
| AW650 | Δ*rlpA*, Δ*mltC* | This study |
| AW656 | Δ*rlpA*, Δ*mltC*, Δ*mltE<>FRT-cat-FRT* | This study |
