## Supplemental Figure Legends for "Lytic transglycosylases RlpA and MltC assist in *Vibrio cholerae* daughter cell separation"

**Fig S1. Δ6 LTG exhibits a mild morphological defect and *mltG*$^{-}$*, rlpA*$^{-}$ exhibit no plating defects*.***

**A)** Cultures of WT and Δ6 LTG were grown in LB at 37 °C for 5 hrs and imaged on agarose pads. Dimensions of >800 cells were compared within length and within width with a Student's t-test where *** indicates p< 0.0001. **B)** Overnight cultures of WT, *rlpA::*stop, and *mltG::*stop grown in LB at 37°C were spot-plated onto LB and incubated ~18 hrs at 30°C. All data are representative of at least two biological replicates.

**Fig S2**. ***rlpA::*stop Δ*mltC* mutant exhibits motility defect but no growth defect.**

**A)** RlpA was depleted in WT, Δ*mltC*, Δ*mltDAC*, Δ*mltDA*, and Δ*mltDC* backgrounds by placing its native promoter under control of arabinose induction and growing in the absence or presence of arabinose. Cells were imaged on an agarose pad. **B)** WT, *rlpA::*stop, and Δ*mltC* were grown to ~$OD_{600}$0.5 in M9+0.2%Glu or LB at 37°C and imaged on agarose pads. **C)** Autolysin mutants were grown in 200 µL LB, monitoring $OD_{600}$. **D)** Overnight cultures of autolysin mutants grown in LB were stabbed into 0.3% agarose LB plates and incubated at 30°C for 16 hrs before measuring the diameter of growth. Diameters of mutant strains were compared to WT by a one-way ANOVA and Dunnett's test where *** indicates a p<0.001 and ** a p<0.01. Error bars represent SEM of 3 independent replicates. Scale bars = 5 µm.

**Fig S3. *E. coli* Δ*rlpA* Δ*mltC* Δ*mltE* mutant exhibits slight separation defect. A)** WT, Δ*rlpA* Δ*mltC*, and Δ*rlpA* Δ*mltC* Δ*mltE* were grown at 37°C to ~$OD_{600}$ 0.6 and imaged on agarose pads. Scale bars = 5 µm. **B)** Cells per chain where manually counted (n >100). Gold bar = median.

**Fig S4**. **Resolution of septal autolysin mutant chains is not due to spontaneous suppressors**. Septal autolysin and Δ*rpoS* mutants were grown in LB at 37°C for 24hrs, then back-diluted into LB at 37°C. Cultures were imaged on agarose pads. Scale bars = 5 µm.

**FigS5. Factors secreted during stationary phase do not affect Δ2 LTG chaining defect. A)** WT and Δ2 LTG were grown in the supernatant of an overnight WT LB culture at 37°C and imaged on agarose pads. Cells per chain where manually counted (n >100). Gold bar = median. **B)** Δ*rpoS* and Δ2 LTG Δ*rpoS* were grown in LB at 37°C and imaged on agarose pads and analyzed as in Fig S5A.

**Fig S6. Functional AmiB-mCherry localizes in Δ2 LTG. A)** Expression of $P_{tac}$: *amiB-mCherry* was induced with 1 mM IPTG in a Δ*amiB* background, grown in LB at 37°C to ~$OD_{600}$ 0.6, and imaged on agarose pads. Cells per chain where manually counted (n >100). Gold bar = median. Gold bars = median. **B)** Expression of $P_{tac}$: *amiB-mCherry* was induced with 1 mM IPTG in a Δ2 LTG background, grown in LB at 37°C to ~$OD_{600}$ 0.6, and imaged on agarose pads. Scale bars = 5 µm

**Fig S7. MltE$_{E. coli}$ is not toxic to WT or Δ2 LTG *V. cholerae.*** **A)** Overnight cultures of $P_{tac}$: *mltE$_{E. coli}$* in WT and Δ2 LTG backgrounds were spot-plated onto LB +/- 1 mM IPTG and incubated overnight at 30°C.

**Fig S8. Autolysin-mCherry fusions are functional.** **A)** With the exception of AmiB-mCherry expressed from chromosomal $P_{tac}$, mCherry fusions were expressed from IPTG-inducible vector pHL100mob. Fusion protein expression was induced with 0.2% arabinose or 1 mM IPTG in a WT background grown in LB at 37°C and detected with Genetex polyclonal mCherry antibody. Loading control band detected with BioLegend monoclonal RpoA antibody. **B)** Expression of chromosomal $P_{tac}$: *rlpA-mCherry* or *mltC-mCherry* was induced with 1 mM IPTG in a Δ2 LTG background, grown in LB at 37°C to ~$OD_{600}$ 0.6, and imaged on agarose pads.

**Fig S9. RlpA and MltC localization to the midcell partially depends on OM insertion.** **A)** Outer membrane signal sequences and lipoboxes in LTG amino acid sequences were predicted by the DOLOP algorithm. **B)** WT carrying pHL100 *dsbA$^{ss}$-mCherry-LTGΔss* was grown in M9 + 0.2% glucose/ kan50 at 30 °C, induced with 1mM IPTG after 2hrs, and imaged on agarose pads at $OD_{600}$~0.15. Demographs of *mCherry* fusion localization generated with Oufti. **C)** Expression of $P_{tac}$: *dsbA$^{ss}$-rlpA$_{[18-263]}$* was induced in a WT background with 1 mM IPTG, grown in low salt LB for 4 hrs, and imaged on agarose pads. **D)** Expression of $P_{tac}$: *rlpA-mCherry* was induced in a Δ 2 LTG background, grown in low salt LB for 4 hrs, and imaged on agarose pads.
